# PROTAC-Design-Evaluator (PRODE) : An Advanced Method for in-silico PROTAC design

**DOI:** 10.1101/2023.06.12.544569

**Authors:** A S Ben Geoffrey, Deepak Agrawal, Nagaraj M Kulkarni, Rajappan Vetrivel, Kishan Gurram

## Abstract

PROTAC (proteolysis-targeting chimeras) is a rapidly evolving technology to target undruggable targets. The mechanism by which this happens is when a bifunctional molecule binds to a target protein and also brings in proximity an E3 ubiquitin ligase to trigger ubiquitination and degradation of the target protein. Yet in-silico driven approaches to design these hetero-bifunctional molecules that have the desired functional properties to induce proximity between the target protein and E3 ligase remains to be established. In this paper we present a novel in-silico method for PROTAC design and to demonstrate the validity of our approach. We show that for a BRD4-VHL PROTAC ternary complex known in the literature, we are able to reproduce the PROTAC binding mode, the structure of ternary complex formed therein and the free energy (ΔG) thermodynamics favoring ternary complexation through theoretical computational methodologies. Further, we demonstrate the use of Thermal Titration Molecule Dynamics (TTMD) to differentiate the stability of PROTAC mediated ternary complexes. We employ the proposed methodology to design a PROTAC for a new system of FGFR1-MDM2 to degrade the FGFR1 (Fibroblast growth factor receptor 1) which is overexpressed in cancer. Our work presented here and named as PROTAC-Designer-Evaluator (PRODE) contributes to the growing literature of in-silico approaches to PROTAC design and evaluation by incorporating the latest in-silico methods and demonstrates advancement over previously published PROTAC in-silico literature.

## Introduction

PROTAC is a rapidly emerging technology for target protein degradation yet approaches for in-silico driven PROTAC design remain rather ad hoc and there is a need to establish methods to better rationalize PROTAC design [1-7]. To rationalize PROTAC mediated ternary complex formation, Drummond et.al. (2019) [1] proposed the use of protein-protein docking and search of PROTAC compatible protein-protein docked poses to rationalize PROTAC mediated ternary complex formation. The development in the literature along the protein-protein docking approach to PROTAC ternary complex modelling has recently culminated in the work of Mikhail et.al. (2023) [2] wherein they conclude that the linker compatible protein-protein docked pose corresponding to the Protein of Interest (POI) and the E3 ligase is the most suitable pose favouring the ternary complex formation. Further, co-operativity has been shown by Wenqing, et al (2022) [3] as an important criterion for ternary complex formation. Co-operativity is defined as the ratio of binding constants corresponding to binary and ternary complex formation [3]. Specifically, for the PROTAC mediated ternary complex to be formed, the ΔG_bind_ of binding corresponding to the ternary complex involving the Protein of Interest (POI)-PROTAC-E3 Ligase should be more negative than the ΔG_bind_ of binding corresponding to the binary complexes involving the Protein of Interest (POI)-PROTAC and PROTAC-E3 Ligase [3]. However, an in-silico methodology to accurately predict the PROTAC binding mode and the PROTAC mediated ternary complex structure has not been reported in the literature thus far. While Wenqing, et al have employed ΔG_bind_ calculations retrospectively on already available PROTAC mediated ternary complex structures available in RCSB to rationalize in-silico the thermodynamics and kinetics of PROTAC mediated ternary complex formation, the approach still cannot be applied in practice as the PROTAC mediated ternary complex structure is not known in PROTAC design problems. We have developed here an in-silico methodology to predict the PROTAC binding mode and the ternary complex formed therein mediated by the PROTAC, overcoming this limitation.We demonstrate the ability of the proposed in-silico methodology to reproduce an experimentally known PROTAC binding mode and PROTAC mediated ternary complex structure as in RCSB with PDB ID 8BDX for the BRD4-VHL system.. This allows for ΔG_bind_ calculations developed in the PROTAC literature so far to be used in practice wherein the experimental PROTAC mediated ternary complex structure is not known. Further, the stability of PROTAC mediated ternary complex is determined by the binding constant (K_d_) associated with the ternary. As an additional in-silico component, we also employ Thermal Titration Molecular Dynamics (TTMD) methodology developed by Pavan et.al.[8] to differentiate the binding constants (K_d_) associated with strong and weak binders to PROTAC system to differentiate (K_d_) values of different magnitude associated with a more stable and less stable PROTAC mediated ternary complex. Taken together we believe our method presented in here represents an important contribution to in-silico PROTAC design literature.

Our PROTAC-Designer-Evaluator (PRODE) methodology, as presented in this paper, will help reduce time as well as costs of the PROTAC DMTA cycle and accelerate early stage PROTAC drug discovery.

## Methodology

### Part A: In-silico methodology to reproduce the binding mode of the PROTAC and the PROTAC mediated ternary complex in the BRD4-VHL system corresponding to the PDB ID 8BDX

PROTAC 48 in PDB ID 8BDX tags Bromodomain-Containing Protein 4 (BRD4) with the E3 ligase called von Hippel–Lindau (VHL) for degradation. In our previous work [7], we proposed that the starting point required for in-silico PROTAC design is the knowledge of the binders for the Target Protein of Interest (POI) which is required to be degraded and binder for the E3 ligase found within the localization of the target protein. From PROTAC 48 found in PDB ID 8BDX, we extracted the binder for BRD-4 and the binder for VHL, as well as extracted the linker used to link these two binders. With these starting points of known binders to BRD-4 and VHL we proceed as follows to reproduce the experimentally known binding mode of PROTAC 48 and the structure of the PROTAC mediated Ternary complex.

### Protein-Protein docking to obtain a linker compatible protein-protein pose of BRD4-VHL

The binders of BRD4 and VHL are first docked into their respective targets and the binding mode of the binders associated with BRD4 and VHL is obtained. Next, the proteins BRD4 and VHL along with the binders in the respective binding mode identified are taken for protein-protein docking. The protein-protein docking was conducted using MEGADOCK 4.0 [9] and several top scoring protein-protein docking poses were generated. Next we analyzed the linker compatibility in the poses. Among the top scoring docking poses, the distance between the binders was analysed and the top scoring pose in which the binders to BRD4 and VHL are close enough in proximity to be connected by a linker is taken as the best protein-protein pose for PROTAC design.

### PROTAC design

The distance between the binders in the selected protein-protein docked pose is the key criteria for linker selection. Among the linkers available in PROTAC-DB [10] that match with the distance criteria for the PDB ID 8BDX, we used the linker for PROTAC 48 to connect the binders using custom rdkit.

### PROTAC binding mode identification

For the PROTAC candidate obtained by connecting the binders with the chosen linker, we generated multiple conformers of the PROTAC candidate using a torsional diffusion approach reported by Bowen et.al for conformer generation [11]. The torsional diffusion approach is more efficient than traditional approaches to conformer generation as the torsional landscape of all possible conformers of the PROTAC are sampled more efficiently in the torsional diffusion approach as compared to traditional approaches [11]. The binding mode of the BRD4 and VHL binders in the selected protein-protein docked pose is chosen as the template and an rdkit based alignment of the generated conformers to the template is carried out. The objective is to obtain a conformer with the maximum alignment and minimum RMSD with the template which retains all the interactions present in the binding mode of the binders of BRD4 and VHL and this is expected to be the binding mode associated with the PROTAC candidate.

Once the PROTAC binding mode is identified, the ΔG calculations associated with the binary and ternary complex as proposed by Wenqing, et al. are carried out to understand co-operativity of ternary complexation. A 50-nanosecond molecular dynamics simulation is carried out and a 3-body system is defined for binary and ternary complex MMGBSA calculations. The PROTAC-BRD4 and PROTAC-VHL form the two binary systems while BRD4-PROTAC-VHL and VHL-PROTAC-BRD4 form the two ternary systems. For performing MMGBSA/PBSA [12] ΔG calculations on the trajectories generated from the 50-nanosecond simulation, a receptor and ligand is defined in the binary and ternary system and their associated ΔGs of binding are calculated. For binary ΔG calculations, BRD4 is defined as the receptor and PROTAC the ligand to obtain the ΔG_bind_ of PROTAC-BRD4 and VHL is defined as the receptor and PROTAC the ligand to obtain the ΔG_bind_ of PROTAC-VHL. In the ternary system, BRD4-PROTAC is defined as the receptor and VHL the ligand to obtain the ΔG_bind_ of BRD4-PROTAC-VHL and PROTAC-VHL is defined as the receptor and BRD4 the ligand to obtain the ΔG_bind_ of VHL-PROTAC-BRD4 system. When the ΔG_TER_ of Ternary is lower than the ΔG_BI_, then ternary complexation is favoured according to Wenqing, et al [3], further whether the PROTAC first binds to BRD4 and then forms the ternary by complexing with VHL or whether the PROTAC first binds to VHL and then complexes with BRD4 can be inferred through the ΔG calculations which we carried out.

While PDB ID 8BDX was used as a validation of the proposed in-silico methodology, the proposed methodology was used to design a PROTAC for a novel system. Fibroblast Growth Factor Receptor 1 (FGFR1) was chosen as the Target Protein of Interest and its known binder Erdafitinib was used in the PROTAC design. The appropriate E3 ligase MDM2 was chosen based on literature [13] and its binder Nutlin was used in the PROTAC design.

### Part B: Thermal Titration Molecular Dynamics (TTMD) to differentiate Ternary complexation-ability for PROTAC

Further, PROTAC mediates the formation of the ternary complex with a different K_d_ (binding constant) value. Lower value of K_d_ more stable the PROTAC mediated ternary complex .. Therefore, it is of value to differentiate, in-silico, a relatively low and high K_d_ value in PROTAC DMTA cycle. We adopt methodology developed by Pavan et.al. [8] to differentiate strong and weak small molecule binders for PROTAC system and differentiate the relatively high and low ternary K_d_ (binding constant) value associated with PDB IDs: 8BDX, 8BDT [14] as a proof of concept of the adaptation of TTMD to PROTAC mediated ternary complex system.. TTMD is based on the hypothesis that a weak binder has a less stable binding mode at higher temperature than a strong binder. Therefore, a short MD simulation is conducted, and the interaction fingerprints are computed across the last 5 nanoseconds of the simulation and Tanimoto similarity is computed between the interaction fingerprints across the first and the subsequent frames to understand the retainment of original interactions. It is expected that for a more stable PROTAC mediated ternary complex having a lower K_d_ value will retain more of the original interaction. The MD simulation was carried out in GROMACS [15] and the interaction fingerprint was carried out through PLIP [16].

## Results and discussion

### Part A

The first part of the methodology involves reproducing in-silico the binding mode of PROTAC 48 as found in PDB ID 8BDX and implementing protein-protein docking on BRD4-VHL to identify a linker compatible protein-protein docked pose. In Table 1, top poses based on highest Protein-Protein interaction scores and linker compatibility are shown.

**Table 1:**
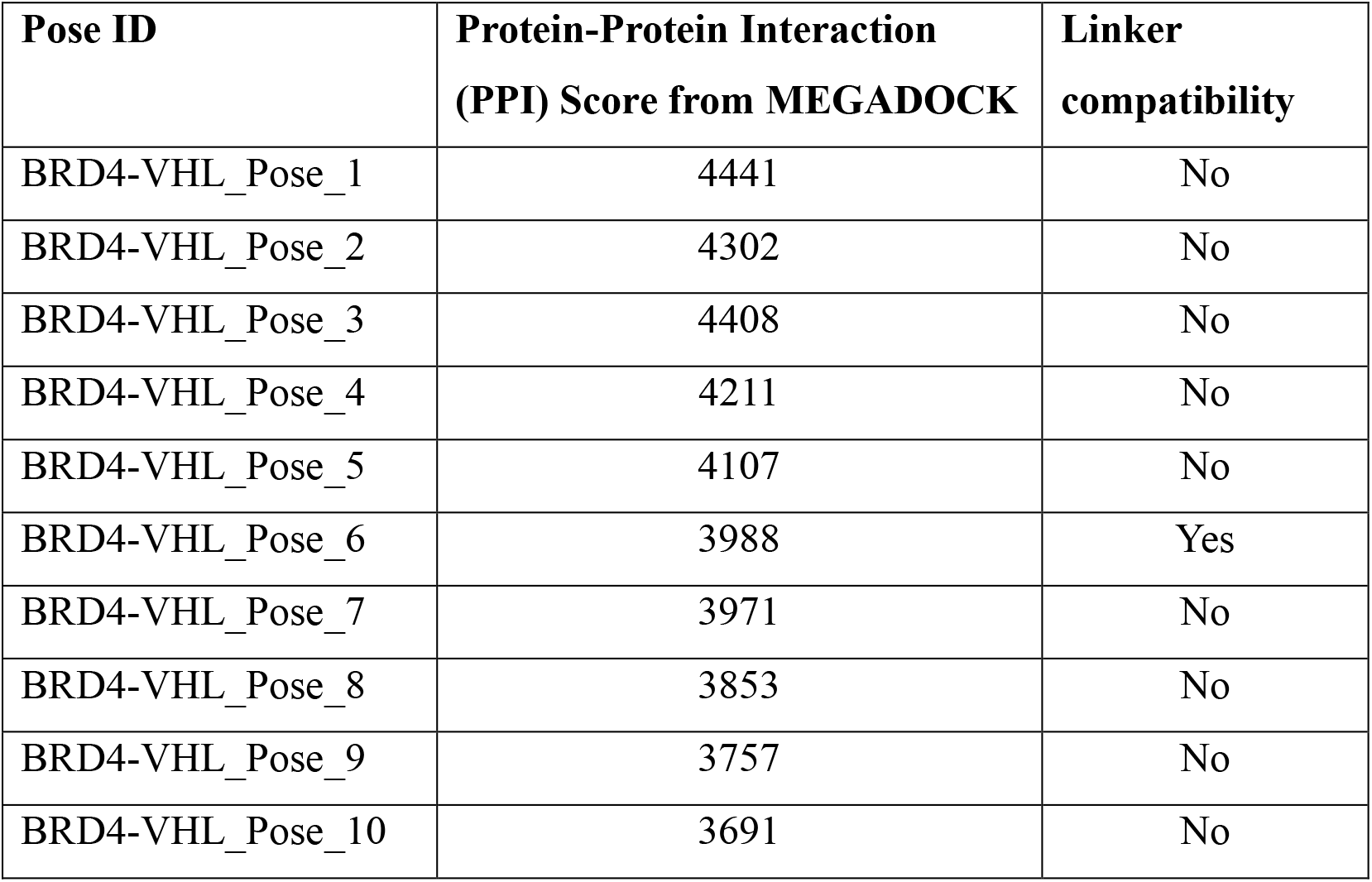
Top scoring poses and linker compatibility.

The linker compatible pose corresponding to Pose ID – ‘BRD4-VHL_Pose_6’ is shown in Figure 1 below. The binding pocket of VHL and BRD4 is orientated along the Protein-Protein Interface which enables the respective binders at the pockets to be connected via a linker making them a linker compatible pose for PROTAC design.

**Figure 1.**
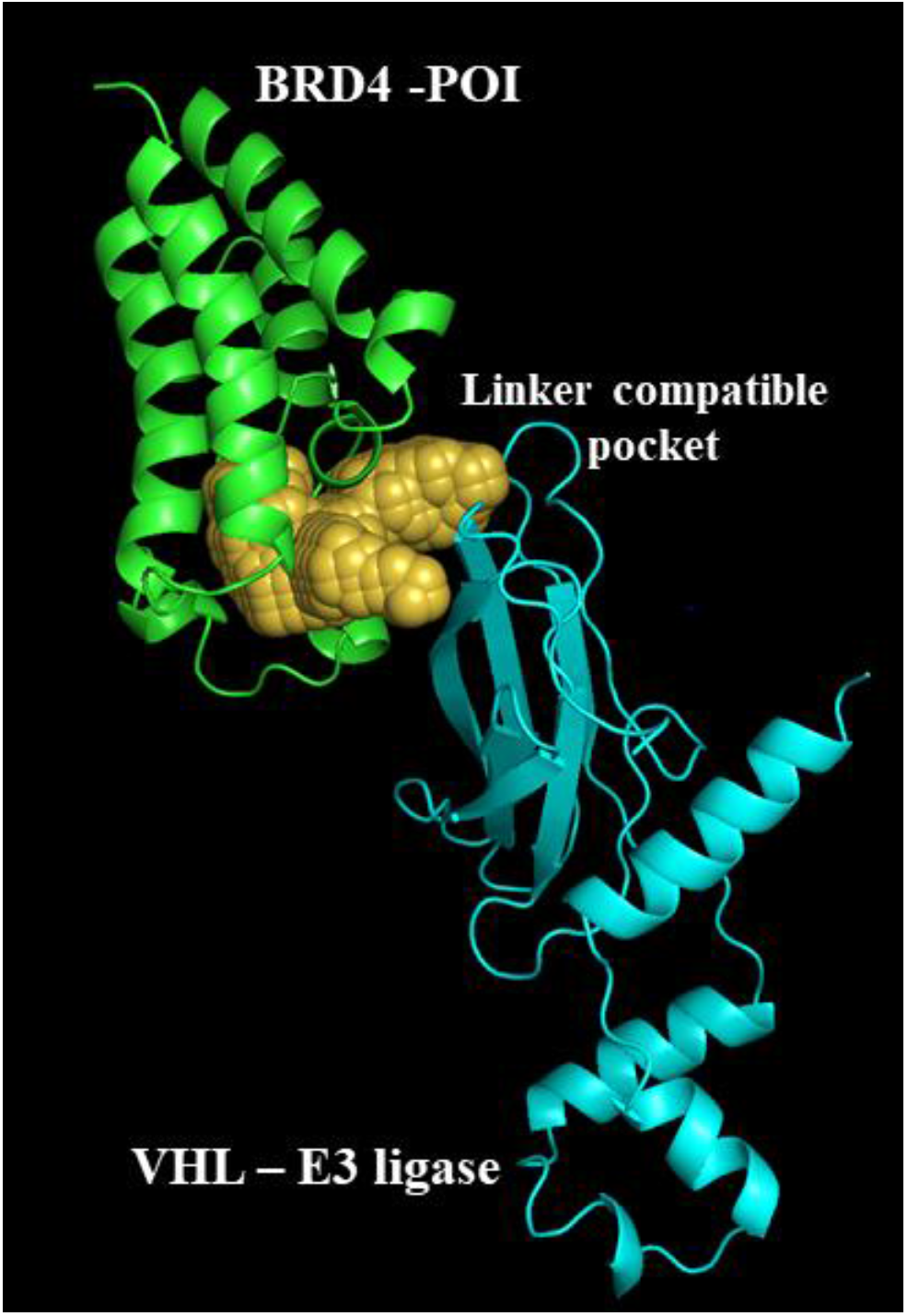
The linker compatible pose corresponding to Pose ID – ‘BRD4-VHL_Pose_6’.

Next step in the methodology to reproduce the binding mode of PROTAC 48 in the BRD4-VHL system corresponding to the PDB ID 8BDX is to reproduce the binding mode of the individual binders in the respective pockets of BRD4 and VHL and the superimposition of the pose obtaining from docking on the crystallographic pose is shown below in Figure 2 for the VHL binder where the crystallographic pose of the binder is depicted in yellow and the docked pose is shown in pink.

**Figure 2:**
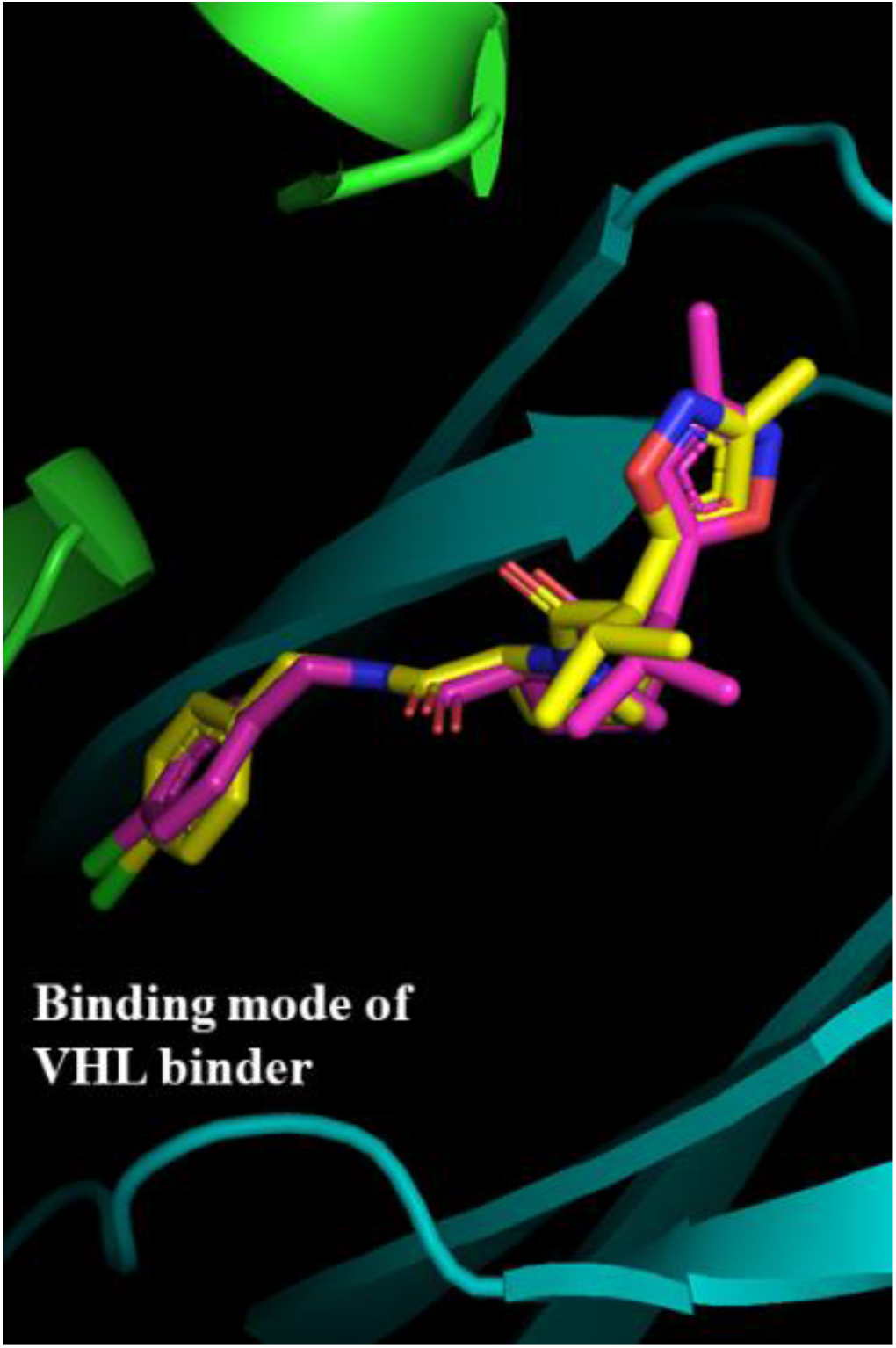
Superimposed docking pose with crystallographic pose for VHL binder.

Similarly, the binding mode of the BRD4 binder was also reproduced and the superimposition of the crystallographic and docked pose obtained is shown below in Figure 3.

**Figure 3:**
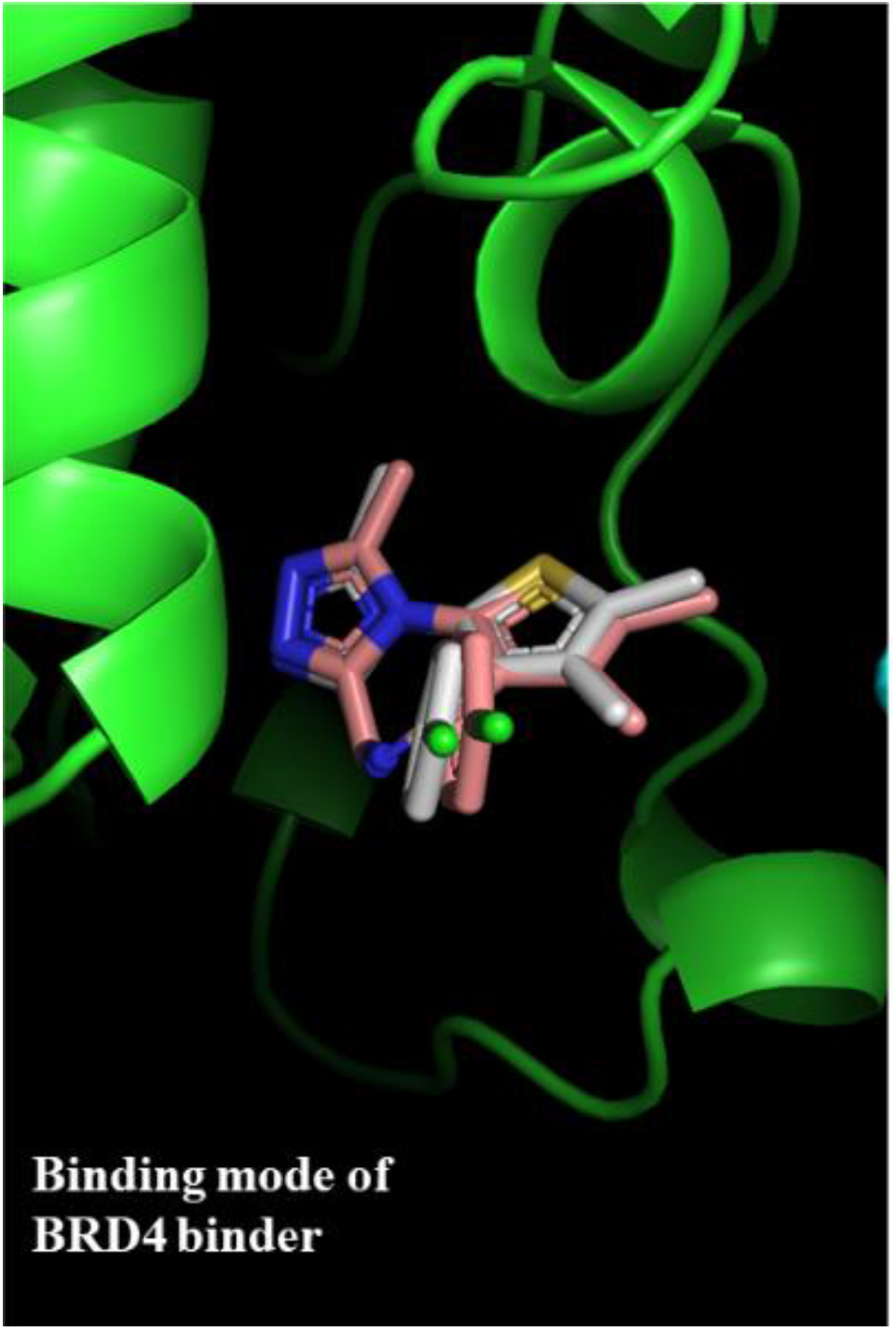
Superimposed docking pose with crystallographic pose for BRD4 binder.

The next step in the proposed methodology to reproduce the binding mode of the PROTAC, is to find a linker of suitable length to link binders of BRD4 and VHL in their respective binding mode such that PROTAC molecule retains all the key interactions present in individual binders of BRD4 and VHL, respectively. For BRD4-VHL system the linker from PDB ID 8BDX was adopted and rdkit was used to connect the binders with the linker and make the PROTAC molecule. 4000 conformers of this PROTAC candidate were generated using torsional diffusion methodology as per Bowen et.al. [11]. The binding mode of the BRD4 and VHL binders individually is taken as template and rdkit was used to obtain the transformation matrix required to align the generated PRTOAC conformers to the template. Transformation matrix encodes the translation and rotations required to align PROTAC conformers to the binding mode of the individual binders of BRD4 and VHL respectively to identify the PROTAC conformer which has maximum alignment to the binding mode of BRD4 and VHL binders and such that PROTAC molecule retains all the interactions present in the individual binders of BRD4 and VHL. The results for the PROTAC conformers with maximum alignment quantified by the RMSD and retainment of interactions quantified by the SF-CNN score is provided in the Table 2 below.

**Table 2.**
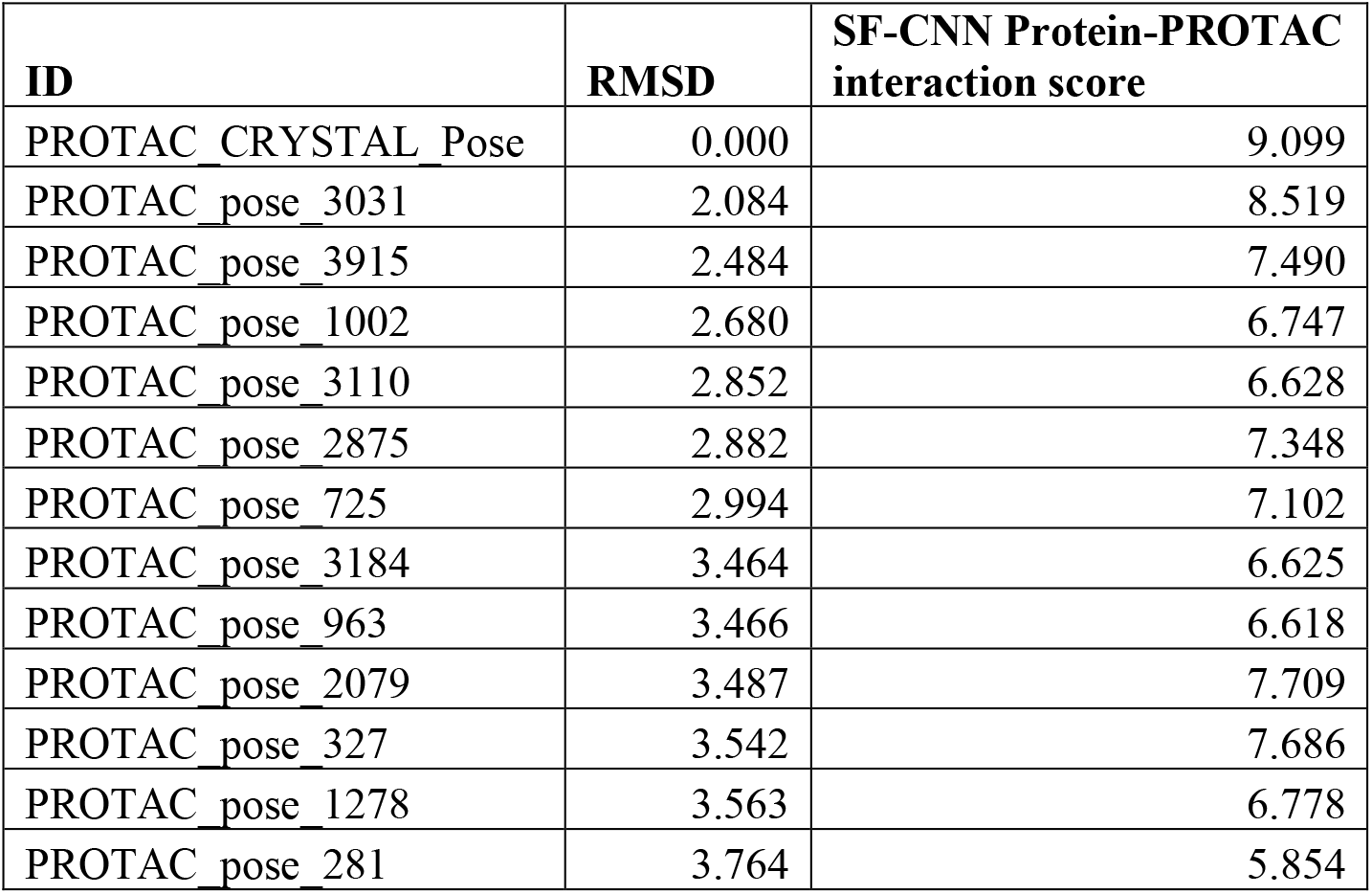
PROTAC conformers with maximum alignment on the basis of RMSD and retainment of interactions.

**Table 3.**
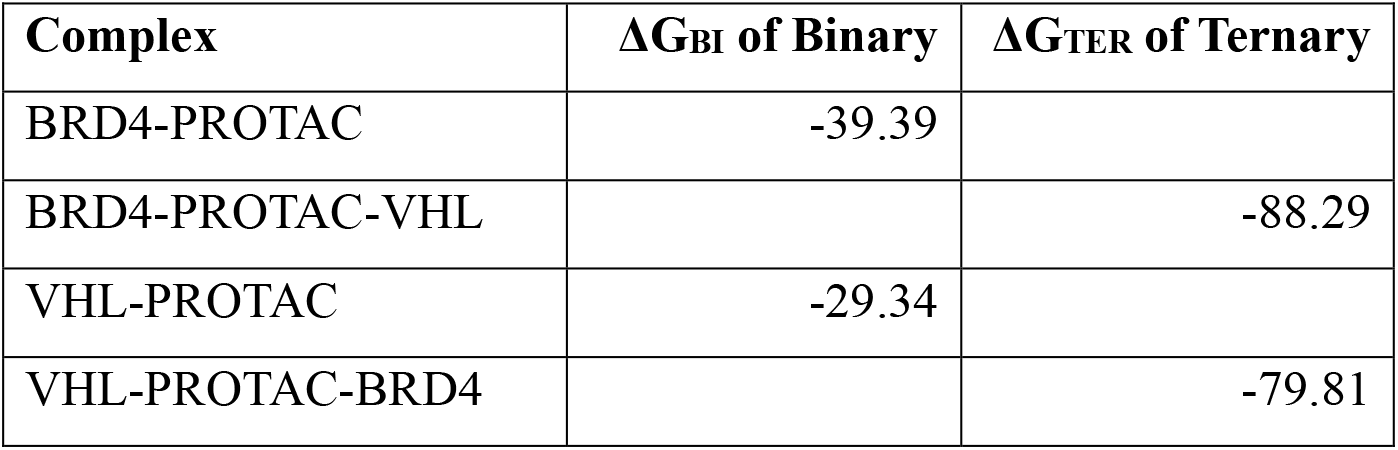
ΔG calculations to rationalize in-silico ternary complex favorability.

The SF-CNN Protein-PROTAC interaction score comes from a deep learning model [17] trained on PDBbind dataset and is meant to provide a quantifiable score to rank interaction between protein residues and small organic ligands. We adopt this model to generate a quantifiable interaction score between the PROTAC molecule and its pocket residues. The first row contains the Crystallographic pose of the PROTAC in PDB 8BDX which obtains a maximum score of 9.099 and among the PROTAC conformers generated through our approach and aligned with the template of the pose of the individual binders of BRD4 and VHL, the 3031^st^ conformer with the PROTAC pose ID, PROTAC_pose_3031 has a maximum alignment with the crystallographic pose of RMSD 2.084 and has a maximum interaction retained as quantified by the SF-CNN Protein-PROTAC interaction score.

The alignment of PROTAC_Pose_3031 with the binding mode of the individual binders of BRD4 and VHL is shown below in Figure 4.

**Figure 4.**
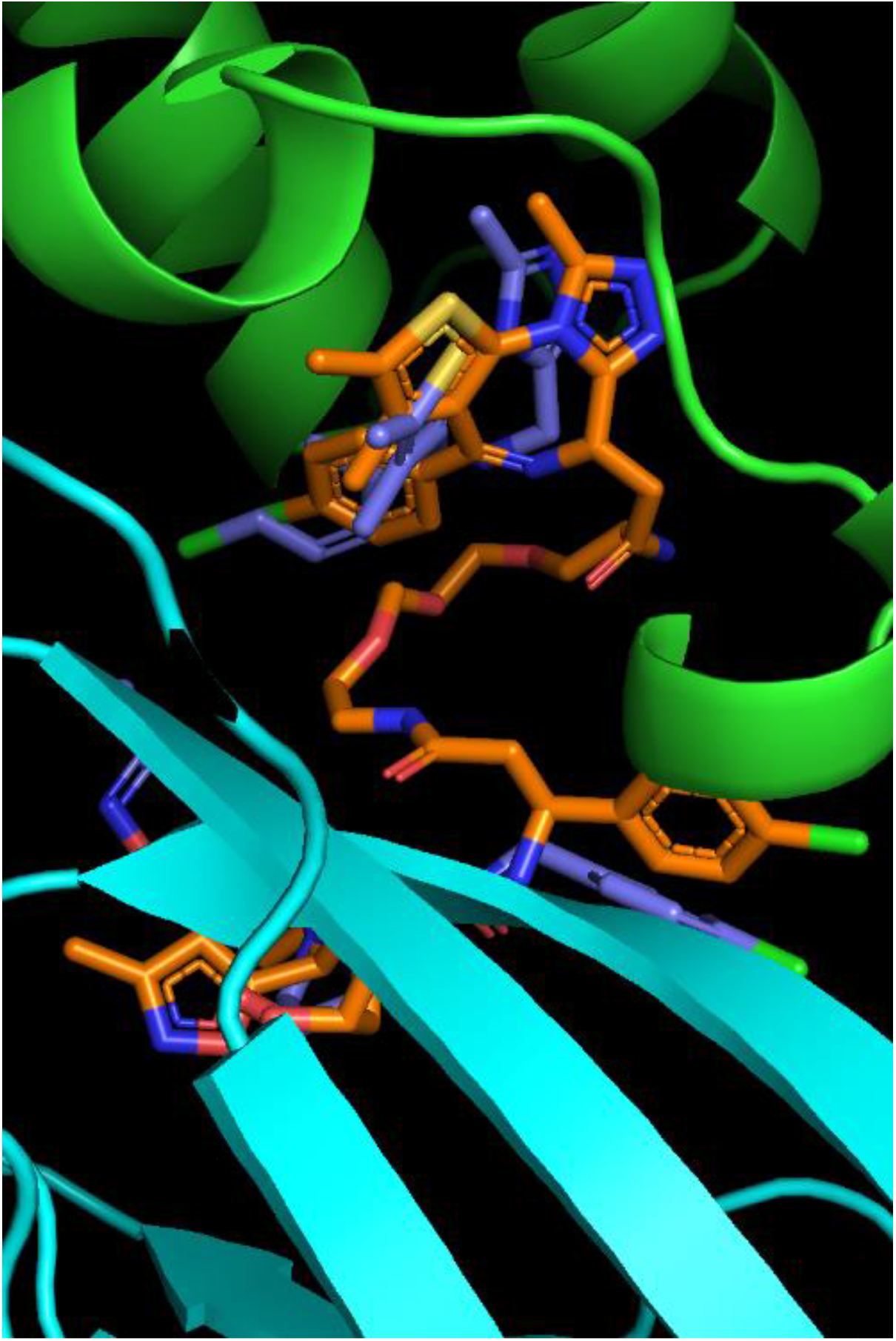
The alignment of PROTAC_Pose_3031 with the binding mode of the individual binders of BRD4 and VHL.

Similarly, the alignment of binding mode of PROTAC_pose_3031 with the crystallographic pose of the PROTAC as found in PDB 8BDX is shown below in Figure 5.

**Figure 5.**
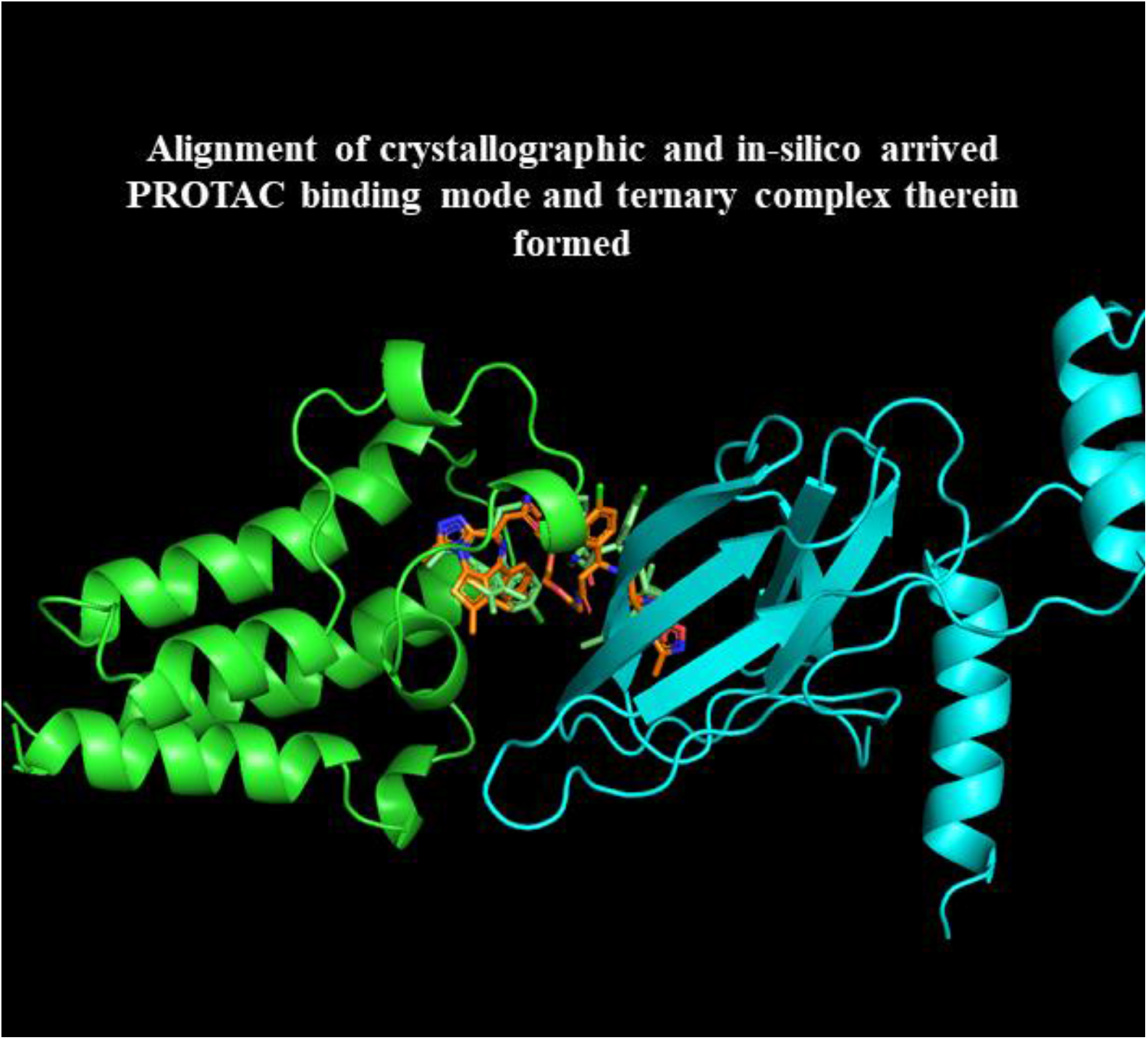
The alignment of PROTAC_Pose_3031 with the crystallographic pose of the PROTAC as found in PDB ID 8BDX.

Further, ΔG calculations as proposed by the Wenqing, et al. were carried out to rationalize in-silico the thermodynamics and kinetics of ternary complex formation.

From this data, it can be inferred that since the ΔG_TER <_ ΔG_BI,_ ternary complexation is thermodynamically favoured.

### PROTAC Design for FGFR1

Following the approach detailed above we next designed a PROTAC for a novel system. Fibroblast Growth Factor Receptor 1 (FGFR1) is overexpressed in cancer and is targeted for inhibition in colorectal cancer. We use the known binder of FGFR1 which is a marketed drug by the name Erdafitinib in the PROTAC design. The E3 ligase Mouse Double Minute 2 (MDM2) was chosen based on literature [13] and its binder Nutlin was used in the PROTAC design.

The structures of Erdafitinib and Nutlin are shown below in Figure 6.

**Figure 6.**
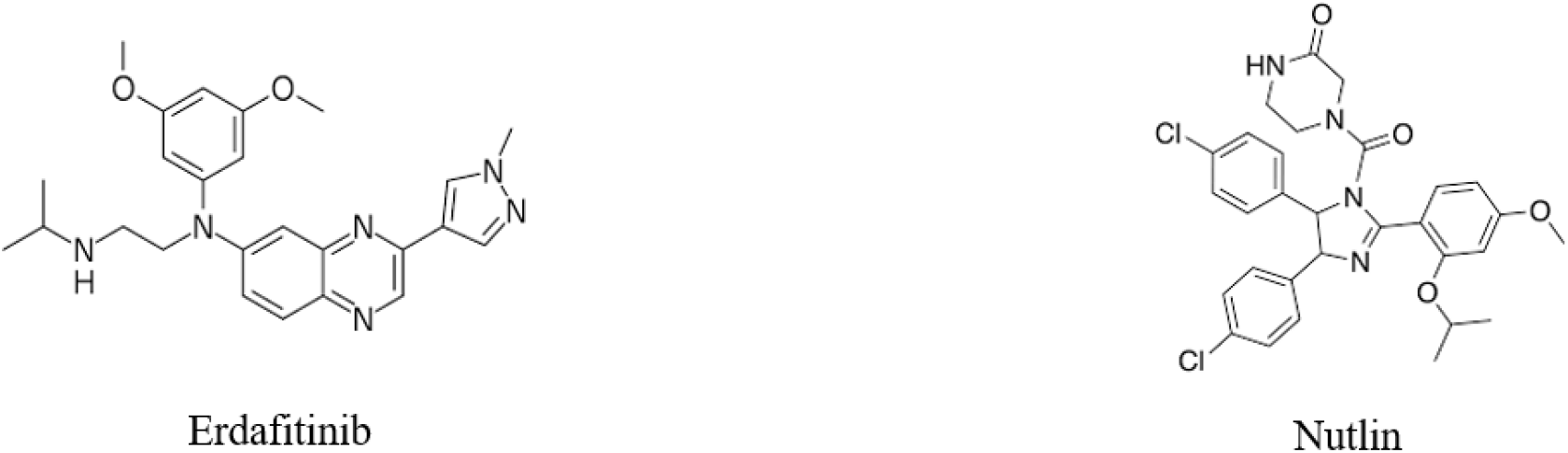
The structures of Erdafitinib and Nutlin.

The approach detailed above was followed to design the PROTAC. The linker compatible protein-protein docked pose between FGFR1-MDM2 was obtained. The individual binding mode of the binders of FGFR1 and MDM2 was obtained. 13 Angstrom was the distance between the binders in their respective binding mode in the linker compatible FGFR1-MDM2 docked pose. Based on the distance needed to link the two binders in their respective binding mode, 4 PEG linkers of length varying around the 13 Angstrom were chosen with Linker IDs 8, 7, 184 and 373 available in PROTAC-DB with the following hyperlinks:

Liknker_8 - http://cadd.zju.edu.cn/protacdb/compound/dataset=linker&id=8

Linker_7 - http://cadd.zju.edu.cn/protacdb/compound/dataset=linker&id=7

Linker_184 - http://cadd.zju.edu.cn/protacdb/compound/dataset=linker&id=184

Linker_373 - http://cadd.zju.edu.cn/protacdb/compound/dataset=linker&id=373

PROTAC candidates were generated out of the different linkers and 4,000 conformers were generated for each PROTAC candidate and were aligned to the template of binders of FGFR1 and MDM2 in their individual binding mode to check for the PROTAC candidate with maximum alignment indicated by a minimum RMSD and maximum retained of interactions present in the binding mode of FGFR1 and MDM2 binders. It was found that the PROTAC candidate made with Linker_184 had minimum RMSD and maximum retainment of interactions. The corresponding superimposed pose of the PROTAC_184 on the binders of FGFR1 and MDM2 is shown below in Figure 7.

**Figure 7.**
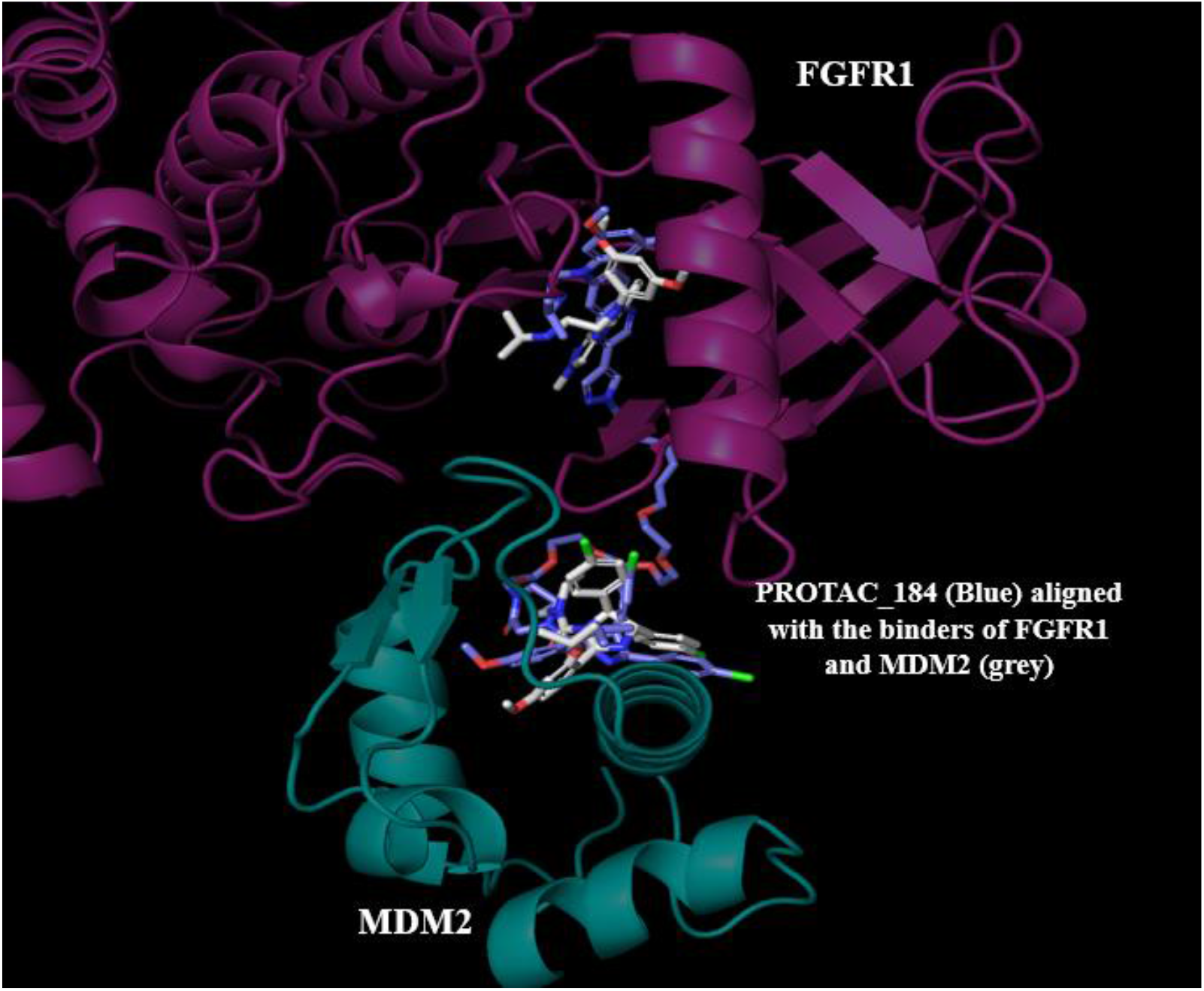
Superimposed pose of the PROTAC_184 on the binders of FGFR1 and MDM2.

Next, ΔG calculations as proposed by the Wenqing, et al. were carried out to rationalize in-silico ternary complex thermodynamic favourability and the results are tabulated below in Table 4.

**Table 4.**
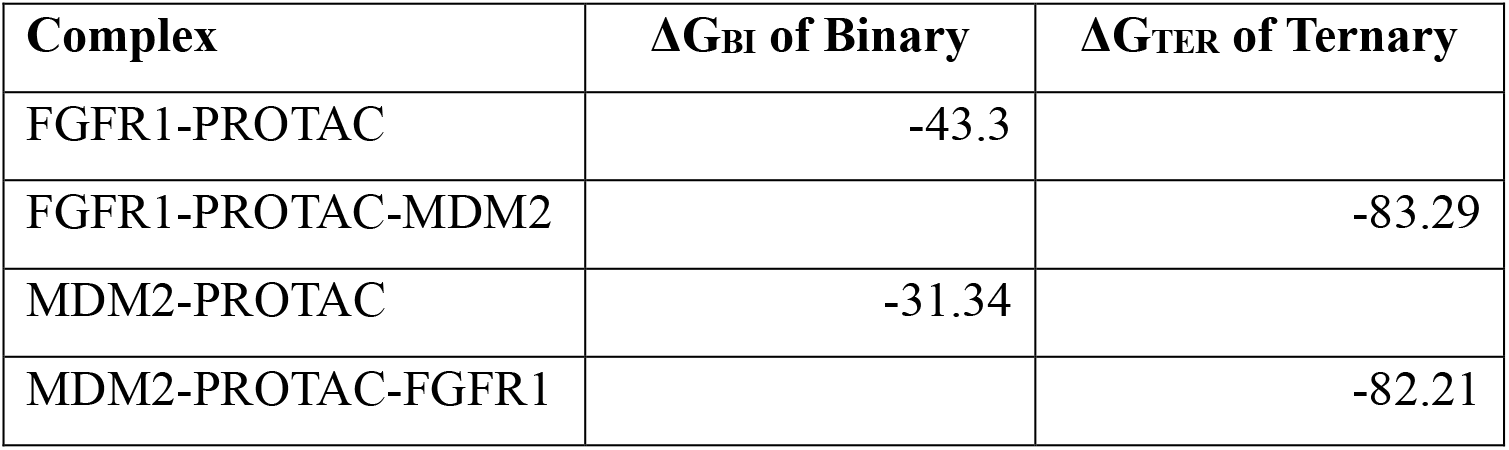
ΔG calculations to rationalize in-silico ternary complex favorability for FGFR1.

From above calculations ternary co-operativity and ternary complexation thermodynamic favorability is concluded positively.

### Part B

In this section we adopt the TTMD methodology developed by Pavan et.al. to differentiate the ternary complex stability mediated by two PROTACs with a relatively higher and lower K_d_ value for the PROTAC mediated ternary complex. We execute a 5-nanosecond classical MD simulation of the PROTAC mediated ternary complex of PDB 8BDX and PDB 8BDT which have two different K_d_ values across a temperature ramp of low (300K) and high (450K). The interaction fingerprints are computed across the generated trajectories and Tanimoto similarity of interaction fingerprints of the first and the subsequent frames of the 5 nanosecond of the simulation is computed to determine the interactions retained. It is expected that a more stable complex will have more interactions retained at a higher temperature than a less stable complex. The results obtained for PDB 8BDX and PDB 8BDT are tabulated below in Table 5.

**Table 5.**
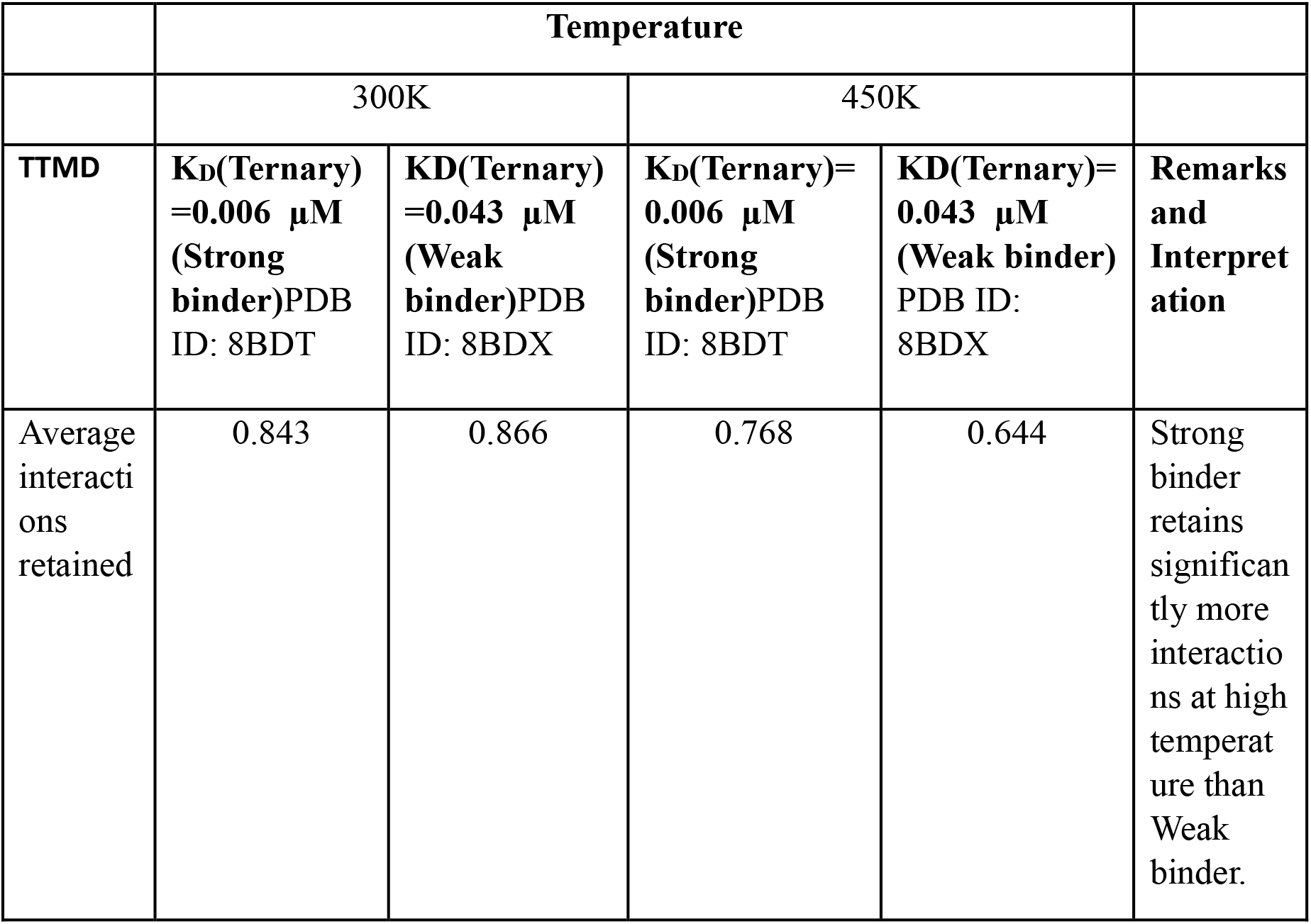
TTMD results for the FGFR1 PROTAC mediated ternary complex.

|  | Temperature |  |  |  |  |
| --- | --- | --- | --- | --- | --- |
|  | 300K |  | 450K |  |  |
| TTMD | $K_D$(Ternary)<br>=0.006 μM<br>(Strong binder)PDB<br>ID: 8BDT | $K_D$(Ternary)<br>=0.043 μM<br>(Weak binder)PDB<br>ID: 8BDX | $K_D$(Ternary)=<br>0.006 μM<br>(Strong binder)PDB<br>ID: 8BDT | $K_D$(Ternary)=<br>0.043 μM<br>(Weak binder)<br>PDB ID:<br>8BDX | Remarks<br>and<br>Interpret<br>ation |
| Average interactions retained | 0.843 | 0.866 | 0.768 | 0.644 | Strong binder retains significantly more interactions at high temperature than Weak binder. |

The average interactions retained is defined as the average of Tanimoto similarity scores calculated between the first and the subsequent frames of the 5-nanosecond simulation.

As it is evident from Table 5 PDB 8BDT which has a lower K_d_ value has more interactions retained at a higher temperature.

To further establish confidence of the use of TTMD method for PROTAC system we carried out the approach for another system taken from literature and found that the trend was repeated for that system. As shown below in Table 6, Strong binder with higher Ternary K_d_ value has higher interactions retained at higher temperature compared to the Weak binder.

**Table 6.**
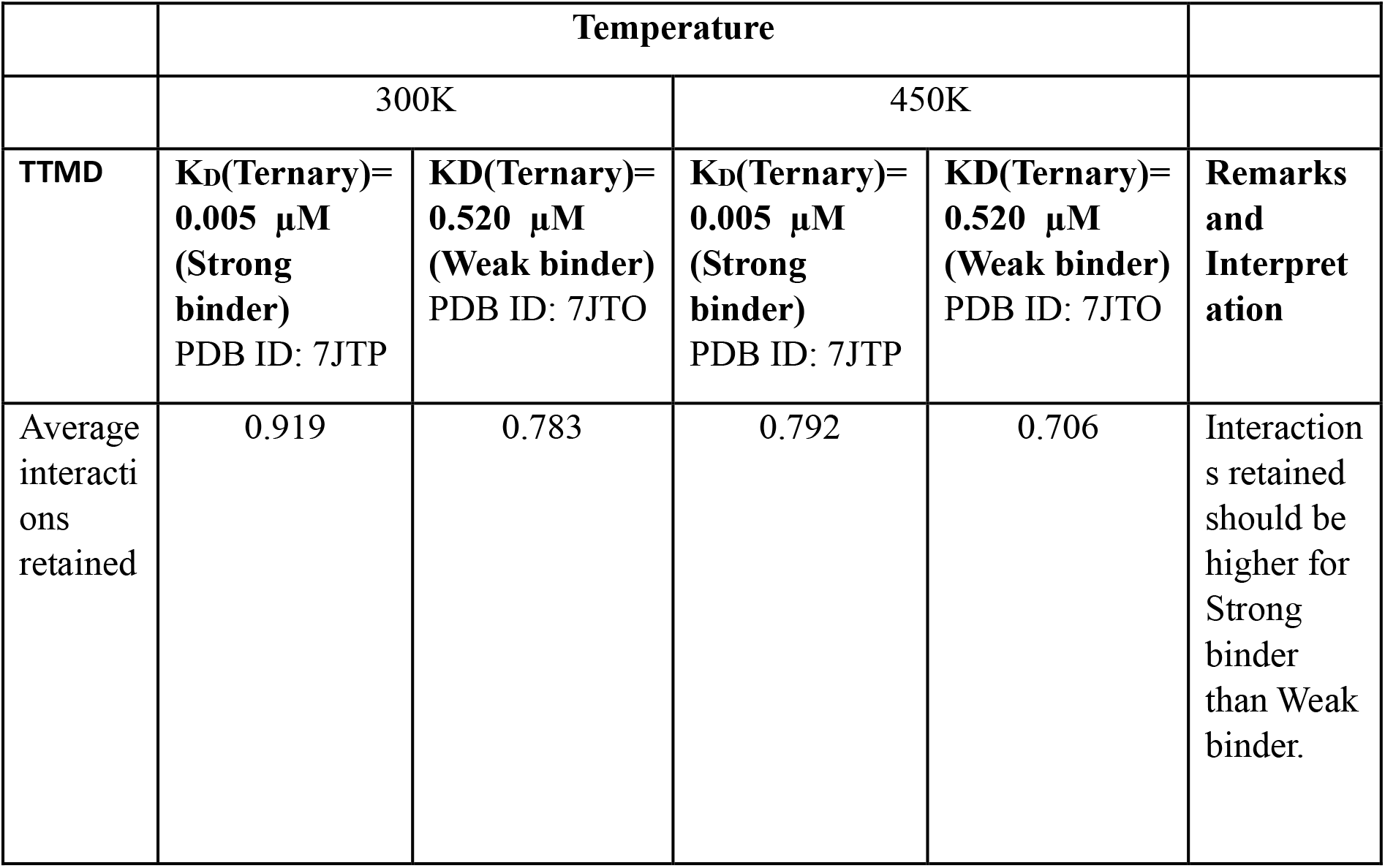
TTMD results for another PROTAC mediated ternary complex.

|  | Temperature |  |  |  |  |
| --- | --- | --- | --- | --- | --- |
|  | 300K |  | 450K |  |  |
| TTMD | **$K_D$(Ternary)=**<br>**0.005 μM**<br>**(Strong binder)**<br>**PDB ID: 7JTP** | **$K_D$(Ternary)=**<br>**0.520 μM**<br>**(Weak binder)**<br>**PDB ID: 7JTO** | **$K_D$(Ternary)=**<br>**0.005 μM**<br>**(Strong binder)**<br>**PDB ID: 7JTP** | **$K_D$(Ternary)=**<br>**0.520 μM**<br>**(Weak binder)**<br>**PDB ID: 7JTO** | **Remarks and Interpretation** |
| Average interactions retained | 0.919 | 0.783 | 0.792 | 0.706 | Interactions retained should be higher for Strong binder than Weak binder. |

Having validated the adaptation for use of TTMD methodology for PROTAC system, we used the TTMD methodology to generate TTMD profiles to understand whether the PROTAC_184 designed for the FGFR1-MDM2 system retained the interactions found in the individual binders of FGFR1 and MDM2 across an increasing temperature ramp. Table 7 below shows TTMD results for FGFR1 binder in PROTAC_184. As we can see the decline in interactions is comparatively small as would be expected of a strong binder.

**Table 7.**
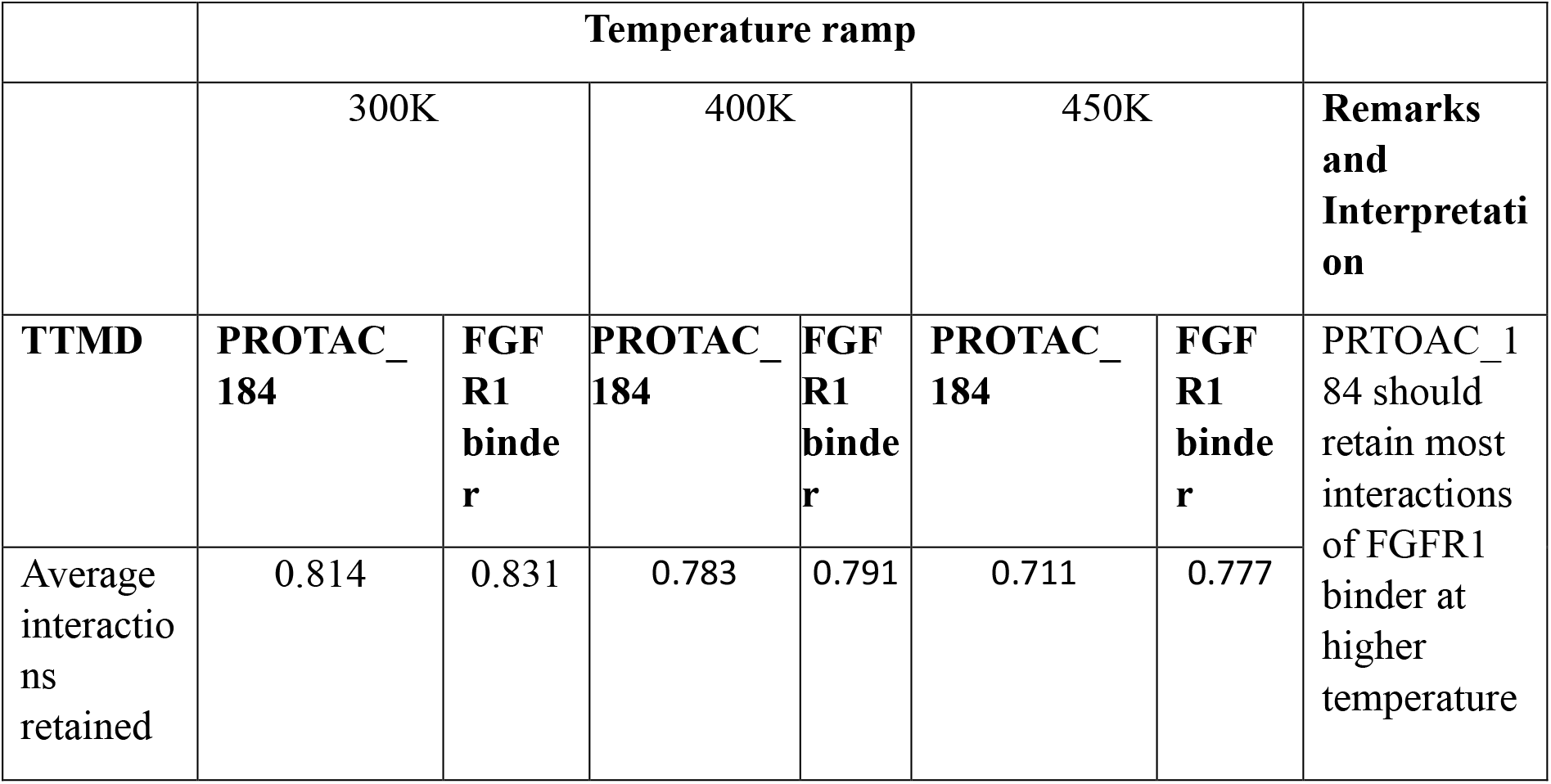
TTMD results for FGFR1 binder in PROTAC_184 mediated ternary complex.

Similarly, Table 8 below shows TTMD results for MDM2 binder in PROTAC_184.

**Table 8.**
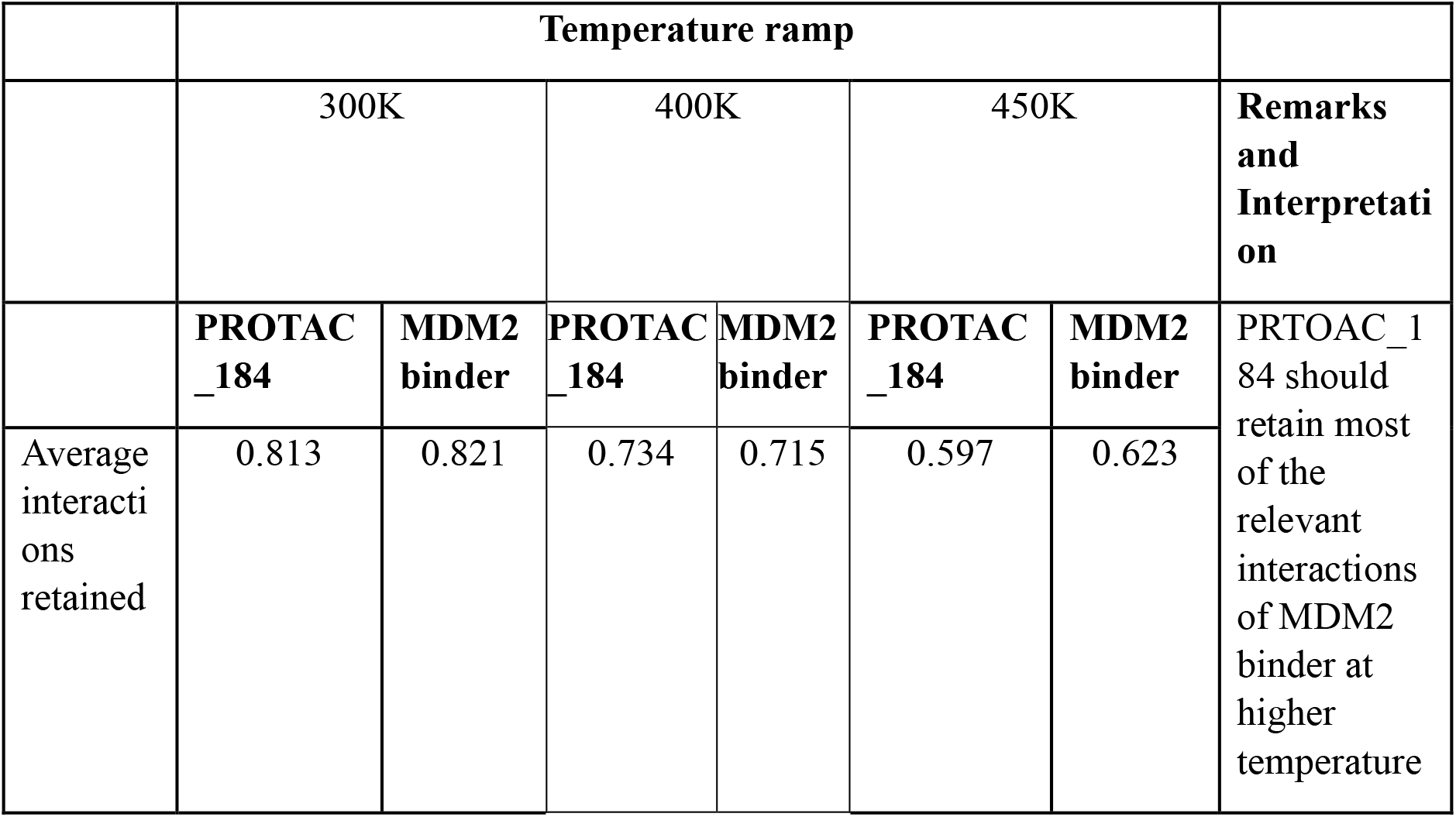
TTMD results for MDM2 binder in PROTAC_184 mediated ternary complex.

From the generated trajectories it is observed that the PROTAC_184 retained the key hydrogen bond interactions (ASP641, ALA564) of the FGFR-1 binder at higher temperatures. Similarly, it was found that PRTOAC_184 retained the hydrophobic interactions (29LEU, 32LEU, 36ILE, 66PHE, 68VAL) of the MDM2 binder at higher temperatures. This is further quantitatively captured in the TTMD profile generated and is indicative of the PROTAC mediating stable ternary complex formation.

## Conclusions

The binding mode of the PROTAC and the ternary complex formed therein as in PDB ID 8BDX was reproduced using the in-silico methodology proposed in our work. The proposed methodology of PROTAC-Designer-Evaluator (PRODE) is as follows. A linker compatible protein-protein docked pose is selected from among the low energy poses obtained from carrying out protein-protein docking between the Protein of Interest (POI) and the E3 ligase. Binding mode of the individual binders of the Protein of Interest and E3 ligase is obtained and based on the distance to link them, linkers are chosen for PROTAC design. A torsional diffusion approach is used to generate the multiple conformers of the PROTAC candidates and are aligned to the individual binders of the Protein of Interest and E3 ligase. A low energy PROTAC conformer with the minimum RMSD and retaining the most interactions as in the individual binders is chosen as the PROTAC binding mode and the resulting ternary complex is used for carrying out further advanced MD based free energy calculations to determine the thermodynamic favorability of ternary complexation. Further, we demonstrate the use of Thermal Titration Molecular Dynamics to differentiate the PROTAC’s ability to mediate a stable ternary complex. The proposed methodology was used to design a PRTOAC candidate for FGFR1 which is overexpressed in cancer and is a target for colorectal cancer.

## Acknowledgements

We are grateful for the various useful discussions we have had with our colleagues at Sravathi AI Technology Private Limited. In particular, we would like to thank Dr. Srinivasan Krishnaswami and Dr. Raghu Bhagavat for their valuable input and insights.

