## Supplementary for "PROTAC-Design-Evaluator (PRODE) : An Advanced Method for in-silico PROTAC design"

| **Pose ID** | **Protein-Protein Interaction (PPI) Score from MEGADOCK** | **Linker compatibility** |
|---|---|---|
| BRD4-VHL_Pose_1 | 4441 | No |
| BRD4-VHL_Pose_2 | 4302 | No |
| BRD4-VHL_Pose_3 | 4408 | No |
| BRD4-VHL_Pose_4 | 4211 | No |
| BRD4-VHL_Pose_5 | 4107 | No |
| BRD4-VHL_Pose_6 | 3988 | Yes |
| BRD4-VHL_Pose_7 | 3971 | No |
| BRD4-VHL_Pose_8 | 3853 | No |
| BRD4-VHL_Pose_9 | 3757 | No |
| BRD4-VHL_Pose_10 | 3691 | No |

**Table 1: Top scoring poses and linker compatibility.**

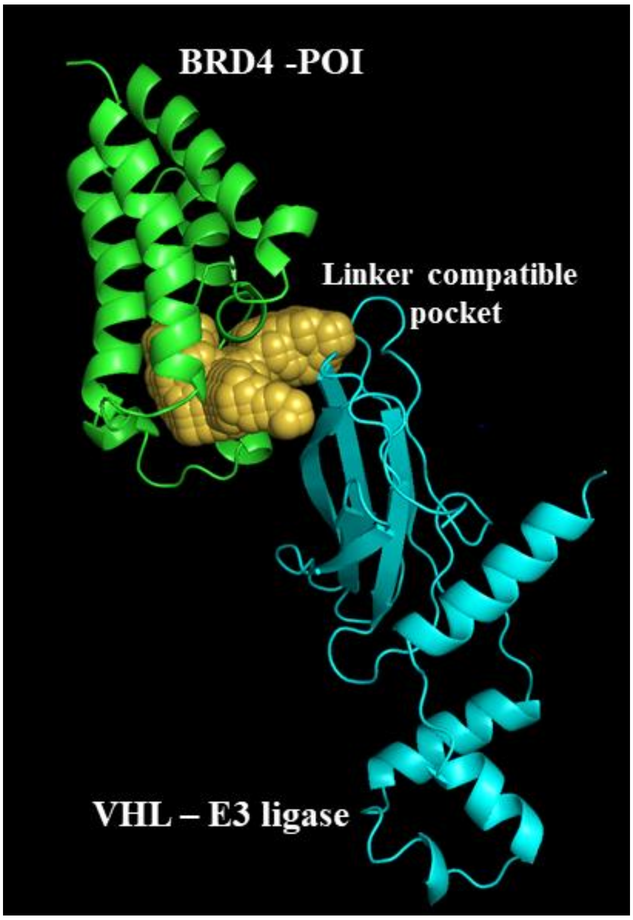


**Figure 1. The linker compatible pose corresponding to Pose ID – 'BRD4-VHL_Pose_6'.**

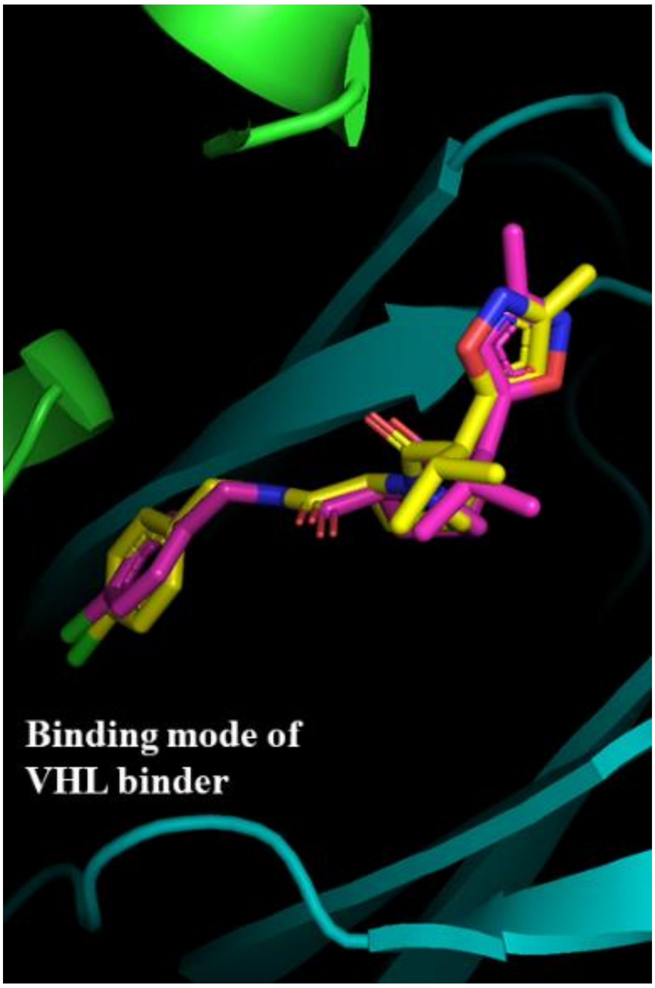


**Figure 2: Superimposed docking pose with crystallographic pose for VHL binder.**

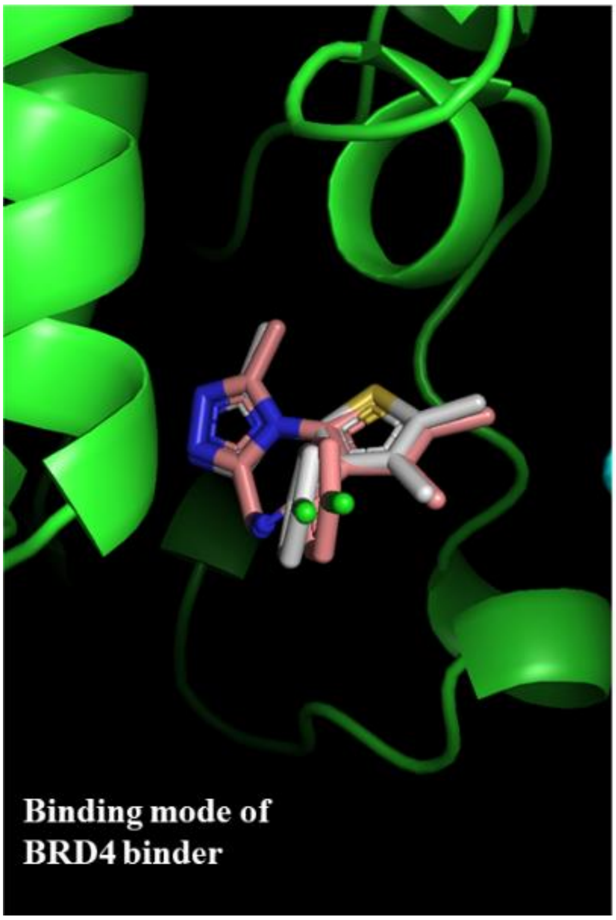


**Figure 3: Superimposed docking pose with crystallographic pose for BRD4 binder.**

| ID | RMSD | SF-CNN Protein-PROTAC interaction score |
|---|---|---|
| PROTAC_CRYSTAL_Pose | 0.000 | 9.099 |
| PROTAC_pose_3031 | 2.084 | 8.519 |
| PROTAC_pose_3915 | 2.484 | 7.490 |
| PROTAC_pose_1002 | 2.680 | 6.747 |
| PROTAC_pose_3110 | 2.852 | 6.628 |
| PROTAC_pose_2875 | 2.882 | 7.348 |
| PROTAC_pose_725 | 2.994 | 7.102 |
| PROTAC_pose_3184 | 3.464 | 6.625 |
| PROTAC_pose_963 | 3.466 | 6.618 |
| PROTAC_pose_2079 | 3.487 | 7.709 |
| PROTAC_pose_327 | 3.542 | 7.686 |
| PROTAC_pose_1278 | 3.563 | 6.778 |
| PROTAC_pose_281 | 3.764 | 5.854 |

**Table 2. PROTAC conformers with maximum alignment on the basis of RMSD and retainment of interactions.**

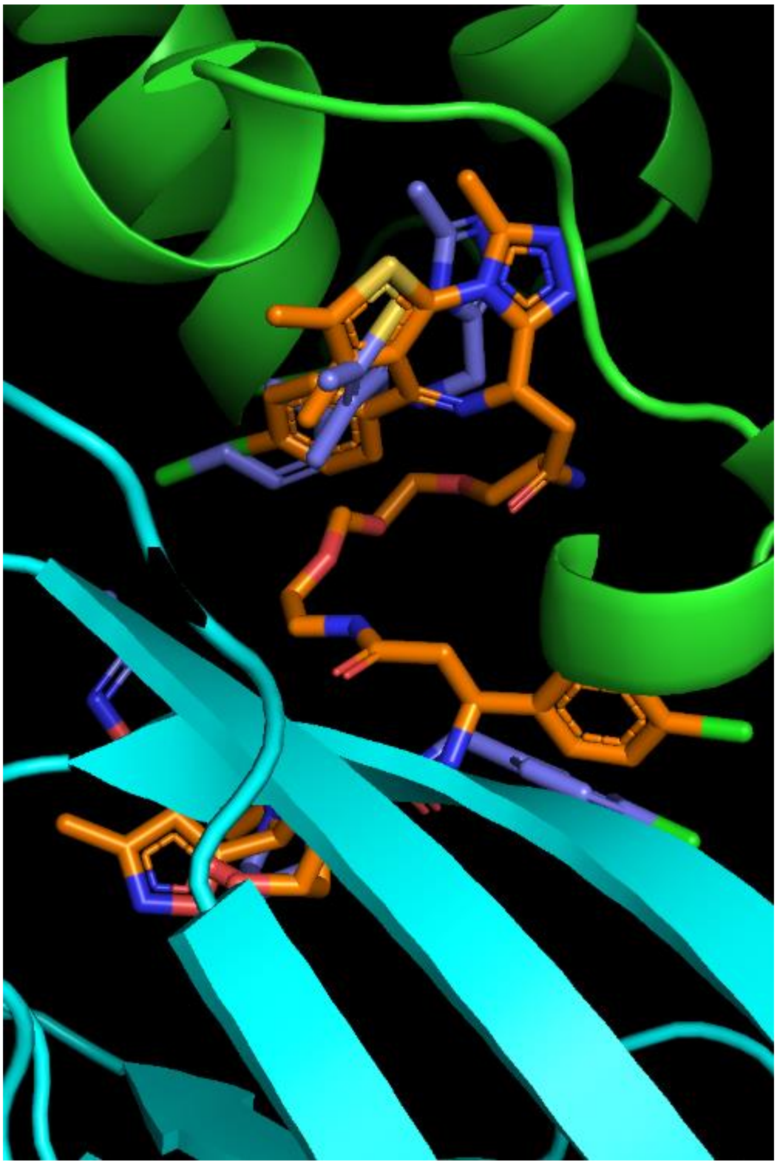

**Figure 4. The alignment of PROTAC_Pose_3031 with the binding mode of the individual binders of BRD4 and VHL.**

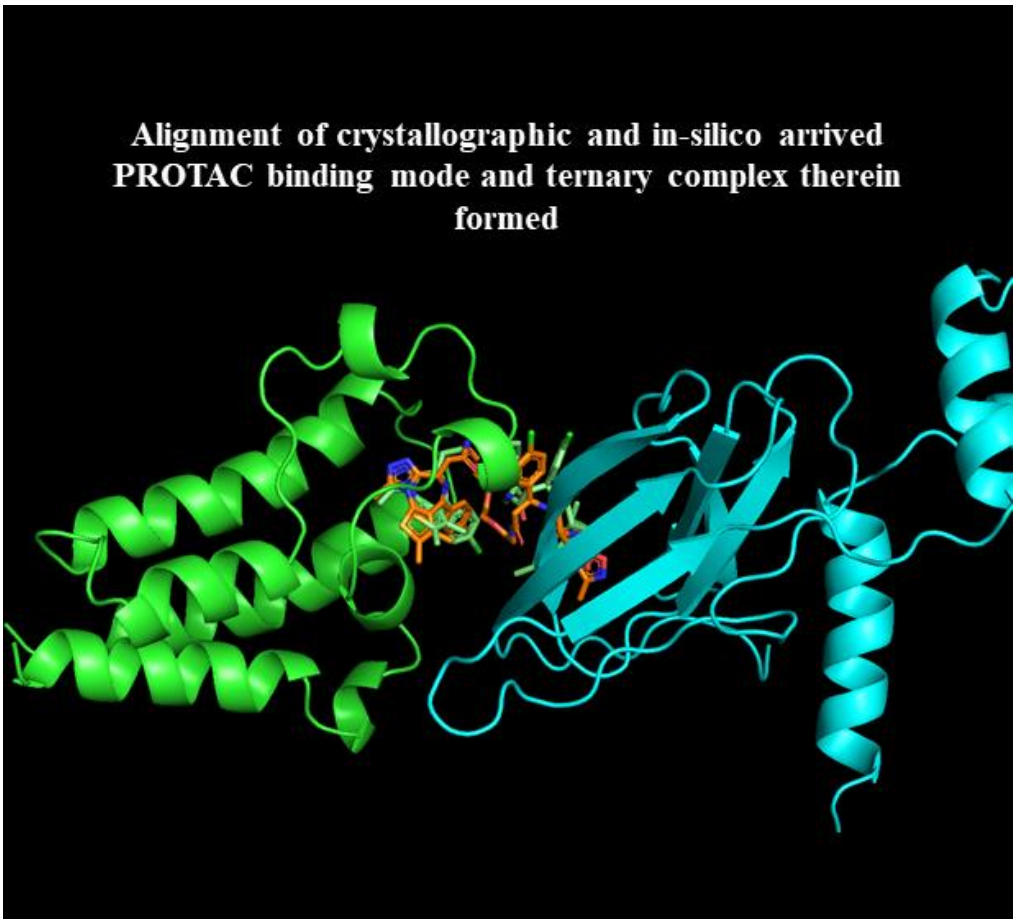


**Figure 5. The alignment of PROTAC_Pose_3031 with the crystallographic pose of the PROTAC as found in PDB ID 8BDX.**

| **Complex** | **$\Delta G_{BI}$ of Binary** | **$\Delta G_{TER}$ of Ternary** |
|---|---|---|
| BRD4-PROTAC | -39.39 | |
| BRD4-PROTAC-VHL | | -88.29 |
| VHL-PROTAC | -29.34 | |
| VHL-PROTAC-BRD4 | | -79.81 |

**Table 3. ΔG calculations to rationalize in-silico ternary complex favorability.**

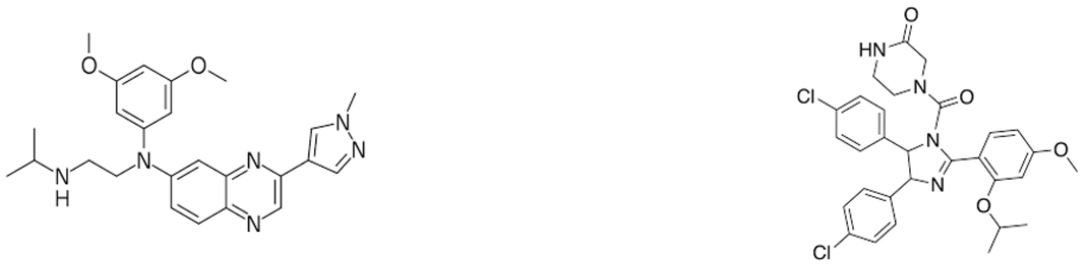

Erdafitinib Nutlin

**Figure 6. The structures of Erdafitinib and Nutlin.**

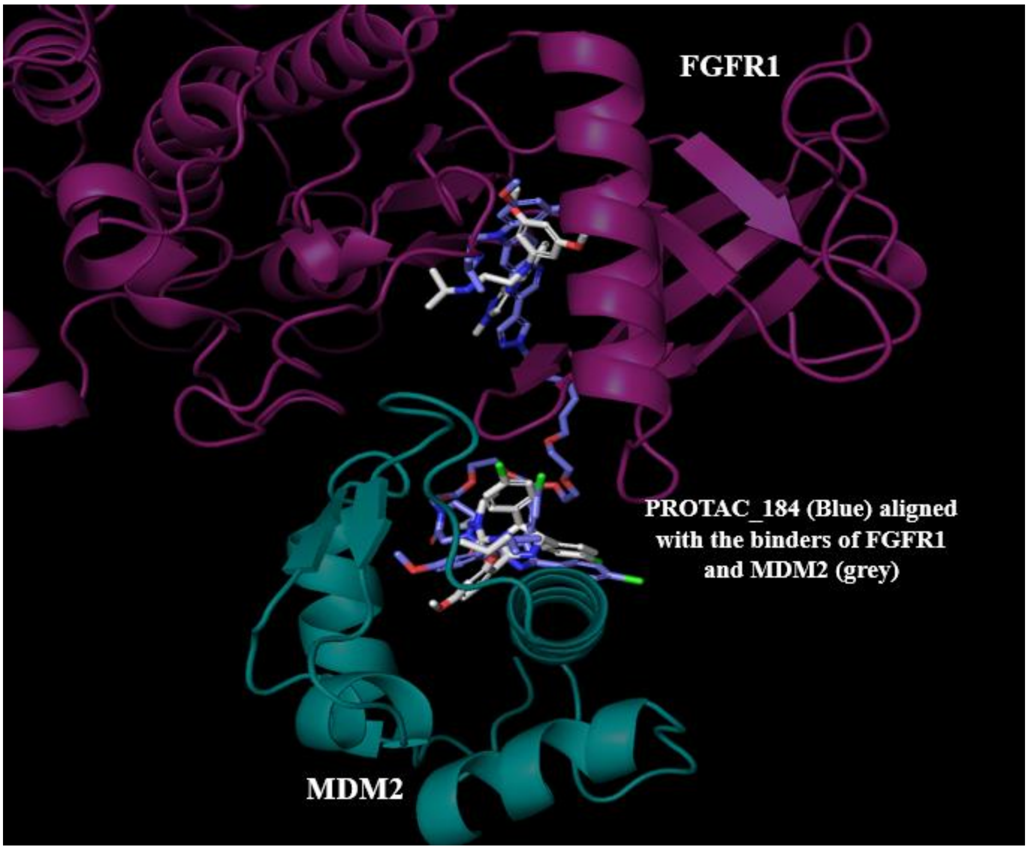


**Figure 7. Superimposed pose of the PROTAC_184 on the binders of FGFR1 and MDM2.**

| Complex | $\Delta G_{BI}$ of Binary | $\Delta G_{TER}$ of Ternary |
|---|---|---|
| FGFR1-PROTAC | -43.3 | |
| FGFR1-PROTAC-MDM2 | | -83.29 |
| MDM2-PROTAC | -31.34 | |
| MDM2-PROTAC-FGFR1 | | -82.21 |

**Table 4. ΔG calculations to rationalize in-silico ternary complex favorability for FGFR1.**

| | **Temperature** | | | | |
|---|---|---|---|---|---|
| | 300K | | 450K | | |
| **TTMD** | **$K_D$(Ternary) =0.006 µM (Strong binder)**PDB ID: 8BDT | **KD(Ternary) =0.043 µM (Weak binder)**PDB ID: 8BDX | **$K_D$(Ternary)= 0.006 µM (Strong binder)**PDB ID: 8BDT | **KD(Ternary)= 0.043 µM (Weak binder)** PDB ID: 8BDX | **Remarks and Interpretation** |
| Average interactions retained | 0.843 | 0.866 | 0.768 | 0.644 | Strong binder retains significantly more interactions at high temperature than Weak binder. |

**Table 5. TTMD results for the FGFR1 PROTAC mediated ternary complex.**

| | Temperature | | | | |
|---|---|---|---|---|---|
| | 300K | | 450K | | |
| **TTMD** | **$K_D$(Ternary)= 0.005 µM (Strong binder)** PDB ID: 7JTP | **KD(Ternary)= 0.520 µM (Weak binder)** PDB ID: 7JTO | **$K_D$(Ternary)= 0.005 µM (Strong binder)** PDB ID: 7JTP | **KD(Ternary)= 0.520 µM (Weak binder)** PDB ID: 7JTO | **Remarks and Interpretation** |
| Average interactions retained | 0.919 | 0.783 | 0.792 | 0.706 | Interactions retained should be higher for Strong binder than Weak binder. |

**Table 6. TTMD results for another PROTAC mediated ternary complex.**

| | Temperature ramp | | | | | | |
|---|---|---|---|---|---|---|---|
| | 300K | | 400K | | 450K | | **Remarks and Interpretation** |
| **TTMD** | **PROTAC_184** | **FGFR1 binder** | **PROTAC_184** | **FGFR1 binder** | **PROTAC_184** | **FGFR1 binder** | PRTOAC_184 should retain most interactions of FGFR1 binder at higher temperature |
| Average interactions retained | 0.814 | 0.831 | 0.783 | 0.791 | 0.711 | 0.777 | |

**Table 7. TTMD results for FGFR1 binder in PROTAC_184 mediated ternary complex.**

| | Temperature ramp | | | | | | |
|---|---|---|---|---|---|---|---|
| | 300K | | 400K | | 450K | | **Remarks and Interpretation** |
| | **PROTAC_184** | **MDM2 binder** | **PROTAC_184** | **MDM2 binder** | **PROTAC_184** | **MDM2 binder** | PRTOAC_184 should retain most of the relevant interactions of MDM2 binder at higher temperature |
| Average interactions retained | 0.813 | 0.821 | 0.734 | 0.715 | 0.597 | 0.623 | |

**Table 8. TTMD results for MDM2 binder in PROTAC_184 mediated ternary complex.**
